# Intracellular niche specialisation drives evolutionary entrapment of endosymbiotic algae

**DOI:** 10.64898/2025.12.22.695913

**Authors:** Irma Vitonytė, Erika M. Hansson, Daniel Malumphy-Montesdeoca, Guy Leonard, Fiona R. Savory, Luke T. Dunning, Thomas A. Richards, Andrew P. Beckerman, Duncan D. Cameron, Michael A. Brockhurst

## Abstract

Endosymbiosis underpins the evolution of complex life and the function of diverse ecosystems, but the mechanisms driving the origin and stability of endosymbiosis remain unclear. Fitness trade-offs between intracellular and extracellular niches could drive the emergence of stable endosymbiosis if adaptation to the intracellular niche reduces fitness in the extracellular niche, reinforcing endosymbiosis through niche specialisation by the endosymbiont. We tested this hypothesis by quantifying phenotypic divergence between endosymbiotic and free-living populations of a facultative algal endosymbiont along trait axes predicted to contrast between its intracellular and extracellular lifestyles. Consistent with endosymbiont specialisation, we observed strong divergence involving multiple traits between endosymbiotic and free-living populations. Specifically, endosymbionts showed convergent losses of multiple nitrogen metabolism pathways, increased export of maltose and glucose, increased sensitivity to acidification, higher and less variable photosynthetic efficiency, and reduced free-living growth rate. Intracellular environments are stable, safe from natural enemies, and come with a reliable supply of specific nutrients, but are acidic and place demands upon endosymbionts to provision their hosts; our data suggest that adapting to this niche is likely to reduce free-living growth and survival of endosymbionts. Our findings support fitness trade-offs between contrasting intracellular and extracellular environments as a mechanism stabilising endosymbiosis by driving evolutionary entrapment of niche-specialist endosymbionts.

## Introduction

Endosymbiosis, where cells of one species (the endosymbiont) live inside the cells of another species (the host), are common across the tree of life [1]. By enabling hosts to rapidly gain new ecological functions, endosymbiosis can drive evolutionary innovation and enhance the functioning of diverse ecosystems [2–6]. However, despite the widespread ecological importance of endosymbioses, their origins and evolutionary stability are poorly understood. Evolutionary theory suggests that explaining the origins of endosymbiosis through mutualism is challenging, because it not only requires that the interaction is beneficial for both parties, but also that the fitness interests of these unrelated species are aligned [7, 8], conditions unlikely to be met for newly formed endosymbioses [9]. Exploitation offers an arguably more plausible route to endosymbiosis [10]: Here, hosts exploit endosymbionts for products or services, and provided hosts can exert sufficient control to enforce the interaction and develop mechanisms for vertical transmission, the fitness interests of the parties progressively become aligned [11]. Such alignment of fitness interests is reinforced where intracellular and extracellular niches impose divergent selection, and fitness trade-offs between these environments drive the evolution of niche specialisation [12]. Given these conditions, as endosymbionts adapt to the intracellular niche, they progressively lose fitness in their ancestral extracellular niche. This process strongly aligns endosymbiont fitness interests with the host’s, such that an initially exploitative interaction now appears mutually beneficial, but entirely through the pursuit of each species’ individual fitness interests [12]. We term this process endosymbiont evolutionary entrapment. Although for most extant endosymbioses such events are lost deep in evolutionary history and unobservable, facultative endosymbiosis, where host periodically sample a new endosymbiont genotype from the free-living population, offer a powerful system in which to study adaptation and divergence between free-living and endosymbiotic niches.

The ciliate *Paramecium bursaria* engages in a facultative endosymbiosis with green algae of the Chlorellaceae. Each host cell houses approximately 100–600 algal endosymbionts arrayed around the interior surface of the host cell membrane, each individually packaged within a perialgal vacuole [13–16]. Hosts, by acidifying the perialgal vacuole [17, 18], provide endosymbionts with amino acids, predicted to be arginine [19] or glutamine [20, 21], from heterotrophy in exchange for photosynthate and elicit algal export of sugars (including maltose). Host cells tightly control the intracellular population size of endosymbionts (termed endosymbiont load) in response to light levels to maximise their benefit:cost ratio: Hosts reduce endosymbiont load both in the dark, where endosymbionts are costly, and in high light, where fewer endosymbionts are required to produce sufficient photosynthate, such that endosymbiont load peaks at intermediate light levels [22, 23]. Hosts achieve this population control through multiple mechanisms, including regulating algal replication [10, 22, 24] and digestion of algal cells [25], with low rates of algal escape in experimental cultures; egestion is uncommon [18, 26]. Given such tight host control, we and others [8, 10, 27] have argued that this symbiosis is founded upon the exploitation of algal endosymbionts by hosts, making it an ideal system in which to test for endosymbiont niche specialisation.

In nature, *P. bursaria* is ecologically widespread in freshwater habitats worldwide where it coexists alongside free-living populations of green algae, enabling hosts to periodically replace their endosymbiont from this free-living pool [28, 29]. It is likely that these intracellular and extracellular niches for the green algae are highly contrasting. For example, hosts provision endosymbionts with organic amino acids [30], but free-living algae primarily acquire inorganic nitrogen compounds, such as nitrate, which are abundant in freshwater environments [21, 31]. Endosymbiotic algae are under selection to release photosynthate upon acidification or be digested, whereas free-living algae experience no such selection [21]. Endosymbiotic algae are taken by their motile hosts to optimal light levels [22] and protected from excess light by algal-produced carotenoids [32, 33], whereas free-living algae are nonmotile and exposed to prevailing conditions.

Due to the facultative nature of this endosymbiosis, host and endosymbiont are separable, meaning that endosymbiotic algae can be isolated and directly compared to their free-living counterparts [19, 22, 34]. A small-scale comparison of 5 endosymbiotic and 4 free-living algal strains found differences in nitrogen and carbon metabolism, with endosymbionts using a narrower range of nutrients than free-living algae [21]. Although consistent with endosymbiont niche specialisation, these strains were taken from strain collections, and as such are not ecologically relevant comparisons for testing the endosymbiont evolutionary entrapment hypothesis.

Here, we tested for divergence between endosymbiotic and free-living algae along multiple trait axes linked to the nutritional environment and host interaction that are likely to vary strongly between intracellular and extracellular niches. We quantified phenotypic divergence across multiple traits for 45 endosymbiotic and 48 free-living closely related *Chlorella*-like algal strains sampled from freshwater bodies located in the northwest of England.

Specifically, we measured autonomous growth rate, photosynthetic efficiency, per capita release of maltose, glucose and trehalose sugars, and use of 3 inorganic and 5 organic nitrogen compounds. We observed strong phenotypic divergence between endosymbiotic and free-living states involving multiple traits. Specifically, endosymbionts had lower autonomous growth rates, higher and less variable photosynthetic efficiency, and higher sugar export and sensitivity to acidification. This was alongside convergent losses of multiple inorganic and organic nitrogen compound metabolism whilst retaining specific amino acid pathways corresponding to those provisioned by hosts. Together, our data provide strong support for the hypothesis that intracellular niche specialisation drives evolutionary entrapment of endosymbiotic algae.

## Results

### Endosymbionts have lower autonomous growth rates than free-living counterparts

We sampled 48 free-living and 45 symbiotic *Chlorella*-like algal strains from 22 freshwater ponds in Northwest England. Algal strains were identified using morphology and SSU rDNA sequences. We constructed a phylogeny using these sequences and sequences from reference strains, confirming phylogenetic groupings for 86 out of the 93 sampled strains (S1 Fig), with the assemblies of the remaining strains being too poor to confidently assign a grouping. As such, all following analyses were additionally tested for robustness using the data subset comprising only the 86 strains from the phylogeny. Symbiotic and free-living lifestyles are intermixed in the phylogeny, although we observed higher SSU rDNA sequence diversity among the free-living strains.

Adaptation to the intracellular niche is predicted to cause reduced fitness of endosymbionts in extracellular environments due to fitness trade-offs [12]. To test this, we compared autonomous growth rates of endosymbiotic and free-living algae in media supplemented with both organic and inorganic nitrogen compounds at pH 7.6 corresponding to the free-living pond environment and pH 5.5 to the acidified peri-algal vacuole. Consistent with our hypothesis, endosymbiotic algae had lower autonomous growth rates than free-living counterparts and particularly so at free-living, neutral pH (Fig 1A; LMM, X^2^ = 112.1, DF = 1, p < 0.001; pH range: 6.13–7.76).

**Fig 1.**
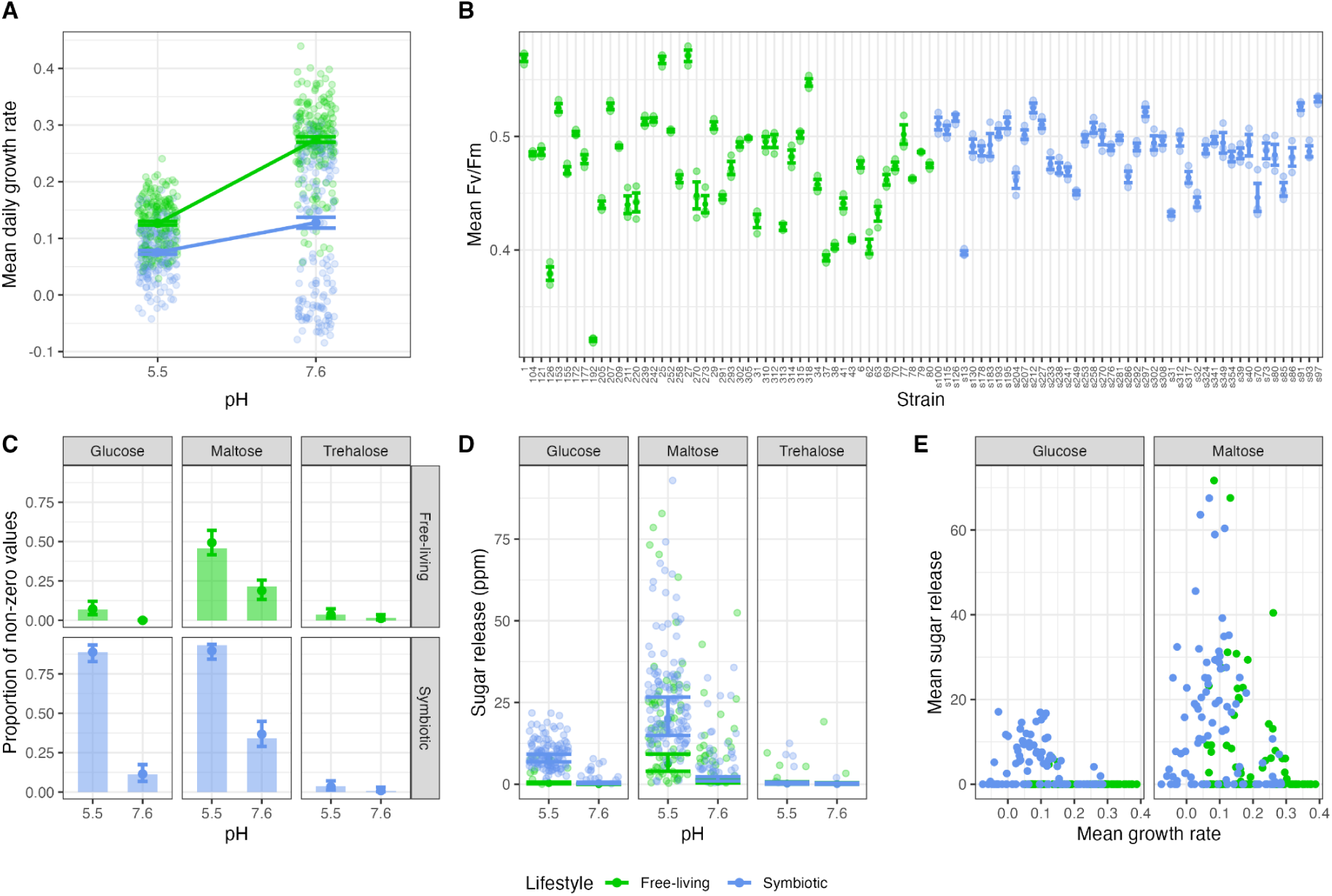
Divergent growth, photosynthetic efficiency and sugar-release phenotypes of free-living (green) and endosymbiotic (blue) strains: A) autonomous growth rate mean with bars for standard error and raw data as transparent points; B) variation in photosynthetic efficiency (Fv/Fm) strain mean with bars for standard error and raw data as transparent points; C) likelihood of sugar release with observed non-zero value proportion as transparent bars and opaque model posterior estimates and 95% CI error bars, D) amount of sugar release when release is > 0 with observed values as transparent jitter and opaque model posterior estimates and 95% CI error bars; E) strain mean sugar release vs strain mean autonomous growth rate.

### Endosymbionts have higher photosynthetic efficiency (Fv/Fm) and export more photosynthate

Endosymbionts are likely to be exposed to more consistent light levels than free-living algae and must export photosynthate upon acidification or be digested [22]. We predicted therefore that endosymbionts should be selected for photosynthetic efficiency and higher rates of sugar export than free-living counterparts. Consistent with this, endosymbionts had higher (LRT, χ²(2) = 6.81, p = 0.033) and less variable (variance ratio: 3.3 FL:S, 95% CI: 2.22–5.01) photosynthetic efficiency (quantified as Fv/Fm; Fig 1B) than free-living algae. (We note that in the robustness test for this data, although the endosymbionts remained less variable, they did not show significantly higher photosynthetic efficiency than free-living strains, although it cannot be ruled out whether this was due to lost statistical power.) Moreover, endosymbionts were both more likely to release sugars and released more maltose and more glucose than free-living algae, and export of both sugars was more strongly elevated under acidic pH in endosymbiont strains (Fig 1C, D; S1 Table). Trehalose export was rare and not associated with lifestyle (Fig 1C, D; S1 Table). Lower autonomous growth rate was associated with higher sugar export for both maltose (Spearman’s rank correlation, ρ = - 0.41, DF = 184, p < 0.001) and glucose (Spearman’s rank correlation, ρ = -0.46, DF = 184, p < 0.001) sugars (Fig 1E), suggesting that such intracellular adaptations are costly.

Together, these data suggest that endosymbionts have adapted to enhance and stabilise photosynthetic efficiency in order to provision more photosynthate to the host, at a cost to their autonomous growth.

### Endosymbionts commonly lose inorganic nitrogen metabolism but convergently retain specific amino acid pathways

Inorganic nitrogen compounds are abundant in extracellular freshwater environments [31], whereas hosts provide endosymbionts with organic nitrogen in the form of amino acids, predicted to be arginine or glutamine [19, 20]. We compared nitrogen compound use between endosymbiotic and free-living algae by measuring growth of all strains in media containing a single organic amino acid (arginine, glutamine, glutamate, asparagine, serine) or inorganic nitrogen compound (ammonium, nitrate, urea). Free-living strains showed higher growth than endosymbiotic strains across all N compounds (Fig 2, S2 Table). Unlike free-living strains, endosymbiotic strains had commonly lost the ability to grow on inorganic N compounds, particularly ammonium and nitrate, and the amino acids asparagine, glutamine and serine. The only N compounds that almost all endosymbiotic strains could use were arginine and glutamate. These data suggest that endosymbionts adapt to the intracellular niche by narrowing their N metabolic niche width, losing pathways for inorganic N compounds and amino acids not provisioned by hosts. Moreover, these data confirm arginine as a likely exchange nutrient [19] but suggest glutamate rather than glutamine [20] as an alternative or additional amino acid used by hosts for endosymbiont provisioning.

**Fig 2.**
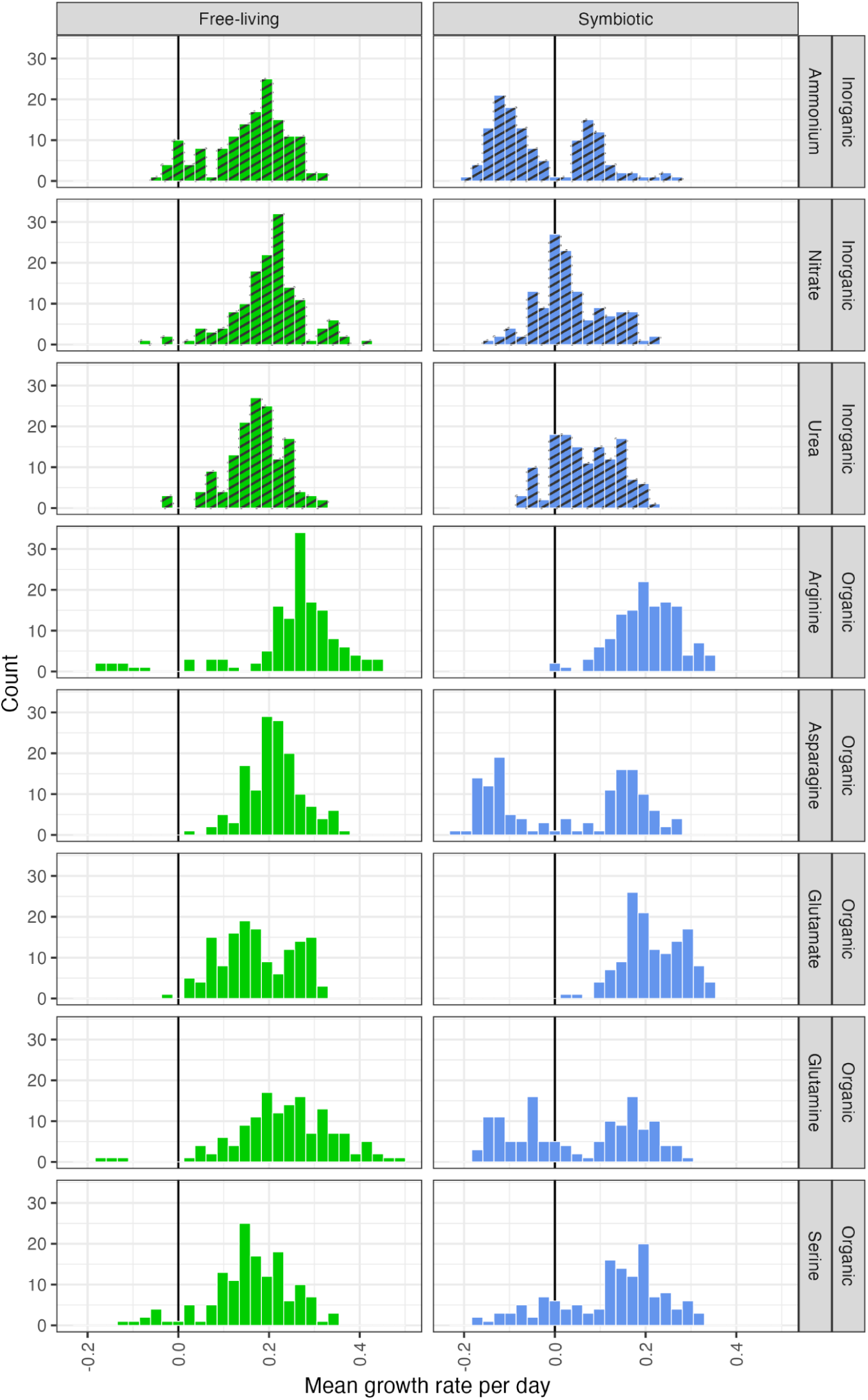
Growth rate distributions of free-living (green) and endosymbiotic (blue) strains on inorganic and organic nitrogen sources. The lefthand column of panels shows free-living strains and the righthand column of panels shows endosymbiotic strains. Rows show different inorganic (hashed bars) and organic (open bars) nitrogen sources (as labelled). Bars indicate counts of strains per growth rate interval.

### Endosymbionts diverge from free-living counterparts along multiple trait axes

Next, we used a Principal Component Analysis (PCA) to reveal multivariate differences among all trait data across strains (Fig 3). Symbiotic and free-living strains occupied distinct clusters (PERMANOVA, R² = 0.10, p = 0.001), with high sugar release and pH responsiveness most strongly loaded toward the symbiotic end, and high autonomous growth rate and high growth rate on most N sources most strongly loaded to the free-living end (S3 Table). This pattern persisted even when the PCA was restricted to phylogroup 1, containing most of the symbiotic strains and their closest free-living relatives (S2 Fig, PERMANOVA, R² = 0.18, p = 0.001), indicating that lifestyle-associated phenotypic structure was not driven by phylogenetic relatedness alone.

**Fig 3.**
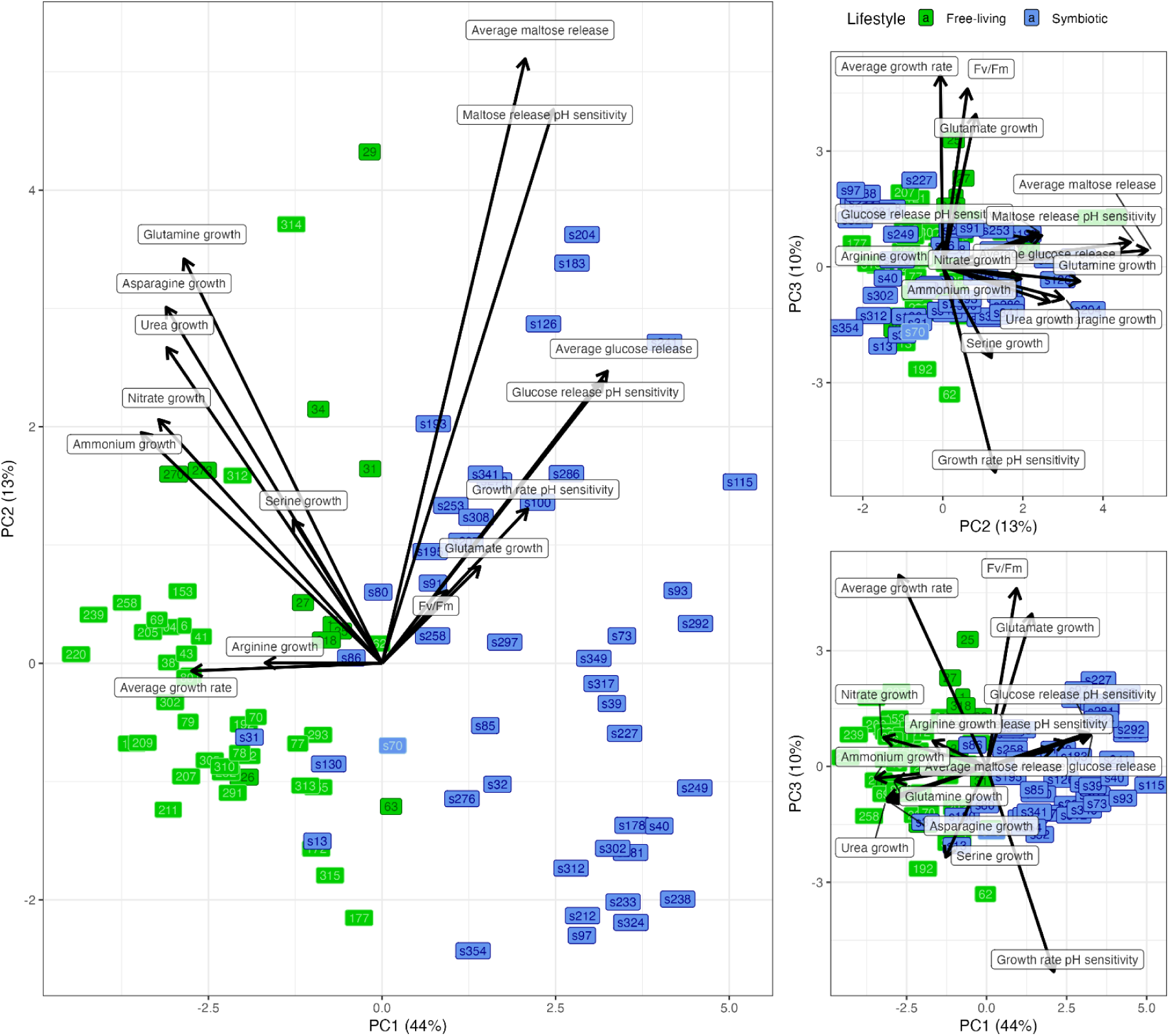
Divergence and clustering of endosymbiotic (blue) and free-living (green) strains along multiple trait axes. Panels show biplots of principal components from a principal component analysis of multivariate trait data. Arrows and labels show loadings of individual variables. Oblongs indicate the position of individual strains in multivariate trait space and are labelled with the strain number; those with darker border/text belong to phylogroup 1 (see Fig S1).

To assess whether inclusion of phylogenetic grouping altered the ordination geometry, we compared the PCA with a Factor Analysis of Mixed Data (FAMD) incorporating phylogroup as a qualitative variable. Individual scores on the first two axes of the PCA and FAMD were tightly correlated (PC1 r = 0.997; PC2 r = 0.985), indicating that including phylogroups does not alter the dominant phenotypic patterns. Although PERMANOVA indicated significant associations between multivariate phenotypic composition and phylogroup (R² = 0.33, p = 0.001) in addition to lifestyle when tested as marginal effects, multivariate dispersion differed among phylogroups (betadisper, p < 0.001), reflecting heterogeneous within-phylogroup phenotypic diversity and unbalanced replication among groups. Taken together, these results indicate an alignment of phylogenetic groups with the existing phenotypic gradients in multivariate space rather than phylogenetic identity driving discrete clustering.

Overlaying pond identity and environmental parameters onto the PCA revealed no consistent association with the ordination axes (S4 Fig). While PERMANOVA detected modest effects of pond identity, nitrate content and pH, these effects were consistently small relative to lifestyle and associated with differences in dispersion (S5 Table), likely owing to uneven lifestyle distribution across ponds. As such, significant pond-level effects should be interpreted as shared pond context and sampling structure, rather than direct environmental filtering, indicating that the observed phenotypic differences between lifestyles are consistent across geographic environments.

Because all traits contributed significantly to multivariate differentiation among lifestyles (S4 Table), a random forest classifier was trained on the same dataset to identify which traits were most distinctive, based on their importance in predicting lifestyle. This model achieved 93.6% overall accuracy (out-of-bag error: 6.45%), with nitrate use, glucose release (both mean and responsiveness to acidification), ammonium use, and autonomous growth contributing most to classification accuracy (S3 Fig). Together these data suggest that selection arising from the intracellular nutritional environment and the symbiotic interaction itself combine to drive divergence of endosymbionts from their free-living counterparts in multiple trait axes.

## Discussion

Endosymbiont evolutionary entrapment occurs when exploited endosymbionts adapt to the intracellular niche, progressively losing fitness in their ancestral extracellular niche due to fitness trade-offs between these contrasting environments [12]. A key prediction arising from this hypothesis is that endosymbiont populations should evolutionarily diverge from the free-living counterparts along trait axes associated with the intracellular environment and host interaction. Using the exploitative facultative symbiosis between *P. bursaria* and *Chlorella*-like algae we show that coexisting populations of endosymbiotic and free-living algae exhibit strong phenotypic divergence. This is strongly evidenced by convergent evolution among multiple endosymbiont strains along multiple trait axes and away from their ancestral free-living state. Endosymbionts narrow their N metabolic niche width to specialise on amino acids provisioned by the host (e.g., arginine), commonly losing ability to grow on inorganic N compounds that are more abundant in freshwater extracellular environments [31]. They also enhance and stabilise their photosynthetic efficiency but export more of their maltose and glucose photosynthate for the host, which comes at a cost to their free-living growth rate.

Together, our data support the hypothesis that intracellular adaptation drives endosymbiont-free living divergence and causes loss of multiple traits important for fitness in the extracellular environment, consistent with endosymbiont evolutionary entrapment through niche specialisation.

Although endosymbiont and free-living algal populations are strongly diverged, there is diversity within both populations and some regions of phenotypic overlap. This is to be expected in an endosymbiosis where hosts recurrently sample new endosymbionts from a free-living pool [28, 29]. Under this regime, the most recently acquired endosymbionts are likely to appear phenotypically free-living-like (e.g., s31, s13, s130 cluster alongside free-living isolates). As such, what we observe in the endosymbiont population is perhaps better conceptualised as an evolutionary trajectory with endosymbiont strains a sample of the population at different stages of their intracellular adaptation. Although *P. bursaria* hosts appear somewhat choosy, preferentially but not exclusively engaging with algae from phylogroup 1, we could even observe this pattern of progressive phenotypic divergence between endosymbiotic and free-living states within the phylogroup.

In endosymbiosis, the algal traits under strongest selection appear to be sugar export and its pH sensitivity. Hosts elicit algal sugar export by acidification of the perialgal vacuole and punish non-exporters by converting the perialgal vacuole into a digestive vacuole and killing the algal cell [13, 18, 25, 35]. Similar host sanctions are a common mechanism by which hosts enforce cooperation from their endosymbionts by policing and punishing cheaters [22]. The strong divergent selection upon sugar export we observe is consistent with the severity of the punishment, here death.

By contrast, N-metabolic niche width is more variable among endosymbionts, although convergent in the N metabolic pathways that are universally retained by all endosymbionts, namely arginine and glutamate. Previous studies have reported that endosymbionts have enhanced amino acid transport systems [30] compared to free-living counterparts [36, 37] and lack nitrate reductase activity limiting utilisation of nitrate [38]. Arginine may be an especially flexible amino acid for provisioning by hosts because it is the only amino acid where uptake is possible across a broad pH range (pH 5.0–6.5) [30]. In contrast, glutamate and asparagine uptake is restricted by external pH, being higher under acidic pH [36, 39]. As such, hosts may use multiple amino acids, perhaps tailoring these to different pH conditions within the perialgal vacuole, to provision and signal to their endosymbionts [27]. These patterns suggest that endosymbionts experience strong and precise selection pressure to retain N metabolic pathways that they need to use the amino acids provisioned by hosts, with other pathways no longer of use. Loss of disused traits can occur due to negative selection if the traits are costly to maintain, or through random genetic drift [40]. Among our strains, we observed closely genetically related free-living and symbiotic strains with divergent N-use phenotypes (e.g., 270, 273, 27 versus s183, s324 have identical SSU rDNA sequences but divergent N use) suggesting that this intracellular adaptation may be rapid and thus likely to be driven by selection on trade-offs.

Near universal retention of arginine and glutamate metabolism in endosymbionts, in contrast to common losses of glutamine metabolism, support previous studies pointing to arginine as the amino acid supplied by hosts [19], but disagree with other studies suggesting that hosts provision glutamine [20]. The evidence for hosts provisioning glutamine is that expression of the host glutamine synthetase gene (glnA), which converts glutamate to glutamine, is elevated in hosts with endosymbionts, and that RNAi knock-down of glnA reduced symbiont load [20]. Notably, these experiments were performed using *P. bursaria* with a different species of algal endosymbiont: *Chlorella variabilis* rather than predominantly *Micractinium* spp. studied here. By contrast, the evidence for hosts provisioning arginine comes from reciprocal pulse chase metabolomics experiments where two divergent *P. bursaria* strains, one with the endosymbiont *M. conductrix* and the other with endosymbiont *C. variabilis*, both showed enrichment within the polyamine pathway downstream of arginine [19]. Together with our data, these patterns suggest that hosts primarily provision arginine regardless of endosymbiont identity but may use other amino acids like glutamate or glutamine for provisioning specific algal species. Testing whether hosts modulate provisioning in response to algal identity requires further experimentation using chemical or reverse genetics

Fitness trade-offs between intracellular and extracellular niches offer a general mechanism stabilising endosymbiosis by driving niche specialisation and evolutionary entrapment of endosymbionts. Here, we show using field isolated facultative endosymbiotic and free-living algae that divergence along multiple trait axes linked to adapting the metabolic intracellular environment and host interactions is associated with loss of free-living metabolic capability and growth. Such mechanisms are unlikely to be unique to this symbiosis given that most eukaryotic intracellular environments will be similar to *P. bursaria’s* and highly distinct from extracellular niches. Consistent with intracellular niche specialisation driven by selection, strong phenotypic divergence was observed even between closely related algae from symbiotic versus free-living niches. Future work will be required to understand the genomic basis of intracellular adaptation and the relative contributions of selection and drift over time in driving evolutionary entrapment.

## Materials and Methods

### Field sites

Twenty-two ponds with confirmed *Paramecium bursaria* populations in the Greater Manchester area were identified and sampled in the summers of 2021 and 2022 (S5 Fig, S6 Table). Physicochemical parameters including pH, conductivity, total dissolved solids and nitrate concentration were measured *in situ*. Ponds ranged from slightly acidic (pH 6.26) to slightly alkaline (pH 7.70), with dissolved mineral content (102–653 µS/cm) and total dissolved solids (51–304 ppm) ranging from low to moderate. The nitrate concentrations did not exceed typical levels of UK waters [41], ranging from 0.44 to 4.30 mg/L.

### Isolation of free-living algal strains

Field samples were taken with a phytoplankton net from the deepest part of each pond. Pond water was gradually filtered (smallest filter pore size 11 µm) to remove larger organisms and left overnight to precipitate at room temperature. Sedimented algae were isolated by siphoning out excess water and then concentrated by centrifuging (2000 RCF, 18C°) for 10 min. The resulting algal pellet was re-suspended in modified artificial WC media (MWC; [42]), streaked on agar plates (MWC with 1.5% agar and ampicillin) and left in our standard growth conditions: 25°C with 50 µE/m^2^/s of light and a 14:10 light: dark cycle. After 1–2 weeks, green colonies were picked and moved into 96-well plates with a liquid MWC media to grow for a further 3–5 days in standard growth conditions. These strains were inspected with microscopy and algae with morphological features corresponding to the *Chlorella*-like genus were chosen for further analysis. Selected strains were re-isolated on agar and moved into modified Bold Basal Media (MBBM, Bold’s Basal Medium with 3-fold Nitrogen and Vitamins; [43]) for maintenance at conditions: 20°C with 33 µE/m^2^/s of light and a 14:10 light: dark cycle. The final collection totalled 48 free-living *Chlorella*-like strains.

### Isolation of symbiotic algal strains

*Paramecium bursaria* were collected with a jug from shoreline of the same ponds. Pond water was filtered gradually (smallest filter pore size 40 µm) to remove larger organisms, and finally a 11 µm filter was used to concentrate ciliates. Isolated *P. bursaria* were pre-adapted to laboratory conditions for two weeks in filtered pond water before moving to modified NCL media [44], where cereal leaf was replaced with 0.25 g/L ground protozoan pellet (CBA053, Blades biological LTD). To isolate endosymbionts, *Paramecium* cells were washed with medium and ampicillin, burst by sonication (20% power for 8 s), streaked on agar plate (MBBM or MWC with amino acids and 1.5% agar with ampicillin) and left in standard growth conditions for 2–4 weeks. Symbiont algal strains were selected and identified the same way as the free-living algae, following re-isolation and maintenance conditions as above. The final collection totalled 45 symbiotic *Chlorella*-like strains.

### DNA extraction and phylogeny construction

To confirm the chosen strains were closely related and fell within the *Chlorella/Micractinium* clade, DNA was extracted for short-read sequencing following the methods used in [45]. Raw Illumina reads were quality-checked and adapted-trimmed with Trim Galore! (default settings) [46] then assembled de novo with SPAdes (default settings) [47].

Using SSU-ALIGN (“DNA” and “Eukarya” mode) [48] we sampled the rDNA gene cluster recovering ∼900 bp of the SSU rRNA gene consistently sampled and directly alignable across 86 of the 93 strains sampled. This target was chosen to enable comparison with previously rDNA-sequenced strains. Overlapping regions of partial SSU rRNA genes were identical across individual and combined assemblies. For each of the 86 strains, one representative partial SSU rRNA sequence was selected, prioritising the longest and then the combined-assembly sequence. All sequences were aligned using Muscle [49] in SeaView [50] and then manually edited and masked.

An initial PhyML [51] tree was built using a GTR+Γ (four rate categories and proportion of invariant sites) substitution matrix, identifying 12 ‘rRNA-types’ (11 highly similar). SSU rDNA sequences from these types were BLASTn-searched against NCBI nr to sample known related taxa. Representative sequences from NCBI, including identical ‘known taxa’ to cover taxonomic affiliation uncertainty and a green-algal outgroup were aligned as above. After refining and masking – excluding the Group I intron reported in some Chlorellaceae [52], which was absent in our strains – the final alignment contained 121 sequences and 777 characters from the front portion of the SSU rRNA region.

The final tree was calculated as above (α shape parameter = 0.67, I = 0.57). The analyses included 100 bootstraps. The resulting tree was arbitrarily rooted on an outgroup of known green algal sequences, and the image was edited in Illustrator. This tree was used to define the phylogroups used in downstream analyses.

### Growth rate in acidic and neutral medium

Green algal growth rate was evaluated in acidic (5.5) and neutral (7.6) pH media, using 100 mM MES monohydrated and MES Na salt for buffering (S7 Table). Algal strains were grown in MBBM without sucrose in standard growth conditions on a shaker (120 RPM) for 7 days prior to the experiment to acclimate and guarantee cells were in the exponential growth phase. Cells were concentrated using a centrifuge (15 min, 2000 RCF), biomass pellets washed with ‘BBM-buffer’ and final biomass concentrated at 10 XG for 4 min. Four replicates of each strain in each testing medium were prepared at a starting density of OD750 0.05 and then placed in a standard growth condition for seven days. The change in cultures growth was assessed with a plate reader (CLARIOstar Plus, BMG LABTECH).

### Photosynthetic efficiency

Algal strains were grown in MBBM in standard growth conditions (S7 Table) on a shaker (120 RPM) for 7 days prior to the experiment to guarantee cells were in the exponential growth phase. The density of algal cultures was evaluated with CytoFLEX (Beckman Coulter) and three technical replicates of 5 × 10^5^ cells/ml concentration of each strain were prepared. Cultures were dark acclimated for 1 hour prior to measurements with LabSTAF fluorometer (Chelsea Technologies), following the manufacturer’s procedure. For maximum quantum yield, measurements were repeated until Fv/Fm stabilized (an average of 200 measurements).

### Sample preparation for screening of sugar release

Maltose, glucose and trehalose secretion was measured in acidic (5.5) and neutral (7.6) pH (S7 Table). Algal strains were grown in MBBM without sucrose in standard growth conditions on a shaker (120 RPM) for 11 days prior to the experiment to acclimate and guarantee cells were in the exponential growth phase. For the experiment, acidic and neutral MBBM media without sucrose and N source was prepared using the same MES buffers as above. Algal cultures were concentrated for 15 min at 2000 RCF, then washed with ‘BBM-buffer’ of pH 7.6 and concentrated again for 10 min at 2000 RCF. Three replicate 2 ml populations of 1 × 10^8^ cells/ml of each strain were prepared in each medium and incubated for 6 hours at standard growth conditions. After incubation all samples were filtered (0.22 µm) and the supernatant frozen (-20°C) for further analysis with ion chromatography.

### Ion Chromatography (IC) of sugar release

The sugar release samples were analysed with DIONEX ICS-6000 SP, AS-AP ion chromatography at Manchester Institute of Biotechnology (MIB, Mass Spectrometry & Separations Facility). The amount of maltose, glucose and trehalose was determined using a CarboPac PA20 Guard Column (30 mm) and a CarboPac PA20 Analytical Column (150 mm). Multi-step gradients of 45 (pH 5.5 medium) and 35 (pH 7.6 medium) min were created to optimise peak separation of all three sugars (S8 Table), using 30 mM NaOH as the mobile phase, with a sample injection volume of 10 µl. Retention time of each sugar in different media are given in S8 Table. Sugar amounts were quantified using 6 concentration levels (STD1, 5, 10, 50, 100, 250) of known standards.

### Growth rate on different organic and inorganic nitrogen (N) sources

Five organic and three inorganic N sources were selected for the experiment with the aim to cover all N metabolism pathways in the study organism and added to MBBM in quantities ensuring an equal number of nitrogen atoms (S7 Table). The medium pH was adjusted to 7.6 and filter-sterilised before the algal strains were added. Algal strains were taken from cultures in exponential growth phase (medium alterations are given in S7 Table) in standard growth conditions on a shaker (100 RPM) and were concentrated with a centrifuge (10 XG for 4 min), then washed with ‘BBM-buffer’ (MBBM medium without sodium nitrate, peptone, sugar or vitamins). The algal pellets were re-suspended with the testing media and left for 18‒20 hrs of medium adaptation. Three replicates of each strain in each testing medium were prepared at a starting density of OD750 0.05 and then placed in a standard growth condition for seven days, when it was measured again with a plate reader (Omega, BMG LABTECH).

### Statistics

All reported test statistics are from analyses carried out on the 93 strains tested, but the robustness of the conclusions were confirmed by rerunning each analysis including only the 86 strains for which SSU rDNA sequences could be used to confirm relatedness. All analyses were carried out using `R` (v4.5.1.). All marginal means for post hoc contrasts were estimated using the `emmeans` package.

Growth rate on different substrates was modelled using a mixed effects model with the `lme4` package, with nitrogen source and lifestyle as fixed effects and strain fit as a random effect with a variable intercept. Significance testing was carried out with `Anova()` from the car package and verified with parametric bootstrapping using the `pbkrtest` package.

Photosynthetic efficiency (average Fv/Fm) was modelled using a mixed effects model fit with the `nlme` package with lifestyle as the fixed effect, split variance by lifestyle and strain fit as a random effect with a variable intercept. Significance testing of the fixed effect was carried out on as a Likelihood Ratio Test (LRT; with ML estimation) against the reduced model.

Sugar release was modelled using Bayesian hurdle gamma regression models with the `brms` package to allow for a two-part structure, the hurdle component modelling the probability of release (i.e., the presence or absence of sugar release), while the conditional component modelled the positive, continuous release values using a gamma distribution with a log link. Each sugar was fit as a separate response variable, with pH and lifestyle as fixed effects for both the hurdle and conditional part of the model. Strain was included as a random effect for the conditional part.

A Principal Component Analysis (PCA) was carried out on the strain means of all measured traits using `prcomp()`. As sugar release as well as autonomous growth was tested in acidic and neutral pH, these were summarised as a mean across pHs to characterise overall performance and difference between pHs to characterise pH sensitivity. Additionally, to confirm inclusion of phylogenetic information did not alter the geometry of the multivariate space, a Factor Analysis of Mixed Data (FADM) was fit using the `FactoMineR` package with phylogroup as a categorical variable alongside the phenotypic data. `adonis2()` as well as `envfit()`, both from the `vegan` package, were used to fit a PERMANOVA to test the effect of lifestyle and phylogroup on the multivariate space as well as the contribution of each variable, and `betadisp()` was used to assess heterogeneity in betadispersion. A random forest classifier was also trained on the dataset using the randomForest package to rank the importance of the variables. Lastly, to test the robustness of our conclusions, a PCA as well as a PERMANOVA testing the effect of lifestyle were carried out on the subset of the phylogroup comprising the majority of the symbiont strains.

## Acknowledgments

This work was supported by grants from NERC (NE/V000128/1) to M.A.B., D.D.C. and A.P.B. and BBSRC (BB/X016439/1) to M.A.B., D.D.C. and T.A.R. We acknowledge David Hopkins for contributions to the fieldwork and lab experimentation.

## Author contributions

M.A.B., D.D.C., A.P. B. designed and conceptualized the project; M.A.B., D.D.C., L.T.D., A.P. B. managed the research project; I.V. field sampling, free-living and symbiotic algal strains isolation, strain identification based on morphological features, maintenance of cultures and performance of experiments; D.M.-M. assisted with measurement of photosynthetic efficiency; I.V. and E.M.H. data analysis; G.L., T.A.R. and F.R.S. performed phylogenetic analysis; M.A.B., I.V., E.M.H. drafted the paper; all authors - contributed to editing and revising the paper.

---

**Fig S1. Phylogeny of 86 strains compared in this project using the front portion of the SSU rRNA encoding gene and demonstrating all strains recovered group closely to known *Micractinium* and *Chlorella* strains.** We note that 41 free-living strains sampled show a higher degree of rDNA variation than the 45 symbiotically isolated strains. The phylogeny was calculated from a masked alignment of 121 sequences and 777 characters using PhyML (REF) with GTR + Γ (with 4 rate categories [α shape parameter = 0.67] 0 and proportion of invariant sites (I = 0.57) correction. 100 bootstrap replicates were conducted.

**Fig S2. Phenotypic divergence of endosymbiotic (blue) and free-living (green) strains belonging to phylogroup 1 in multivariate space with variable loadings.** Panels show biplots of principal components from a principal component analysis of multivariate trait data. Arrows and labels show loadings of individual variables. Oblongs indicate the position of individual strains in multivariate trait space and are labelled with the strain number. All strains belong to phylogroup 1 as defined in Fig S1.

**Fig S3. Phenotypic variables ranked by Mean Decrease Accuracy (importance as predictor) and Mean Decrease Gini (importance for model to be able to make a clean split) for the Random Forest classifier model predicting strain lifestyle.** Where the trait was tested in both neutral and acidic pH, the trait has been summarised as a mean for overall performance and a slope (difference) between performance in the two pHs to indicate pH sensitivity.

**Fig S4. Principal Component Analysis biplots of multivariate trait data overlayed with environmental variables** a) pond pH, b) pond nitrate content, c) pond TDS, d) pond conductivity and e) pond ID.

**Fig S5. Map of pond sites in Greater Manchester area used for fieldwork sampling during June–September in 2021 and 2022.** Map generated using GoogleMaps on 24^th^ July 2025.

**Table S1. Model summary output and selected pairwise contrasts from Bayesian hurdle gamma regression model estimating likelihood of sugar release (hurdle) and amount of sugar released when non-zero (mean).**

**Table S2. Output from mixed effects model testing effect of lifestyle and N source on growth rate along with selected pairwise contrasts.**

**Table S3. Output from envfit analysis showing correlation of phenotypic variables with PCA axes.** The trait was tested in both neutral and acidic pH and has been summarised as a mean for overall performance and a slope (difference) between performance in the two pHs to indicate pH sensitivity.

**Table S4. Results of envfit analysis showing correlations between individual traits and the first two principal components of the multivariate trait ordination.** Vector directions indicate the orientation of increasing trait values in ordination space.

**Table S5. PERMANOVA output testing effects on multivariate centroids of lifestyle interacting with pond identity and associated environmental variables, as well as betadisper output for homogeneity of dispersion tests.** Pond identity and environmental variables were tested in separate models.

**Table S6. Site details, mean values of physicochemical parameters measured *in situ* June–September in 2021 and 2022.**

**Table S7. Alterations of MBBM medium made for algal biomass growth and experiment of each assay.**

**Table S8. Methods of Multi-Step Gradient for sugar release detection in different pH media.**

