## Supplementary Figures for "Intracellular niche specialisation drives evolutionary entrapment of endosymbiotic algae"

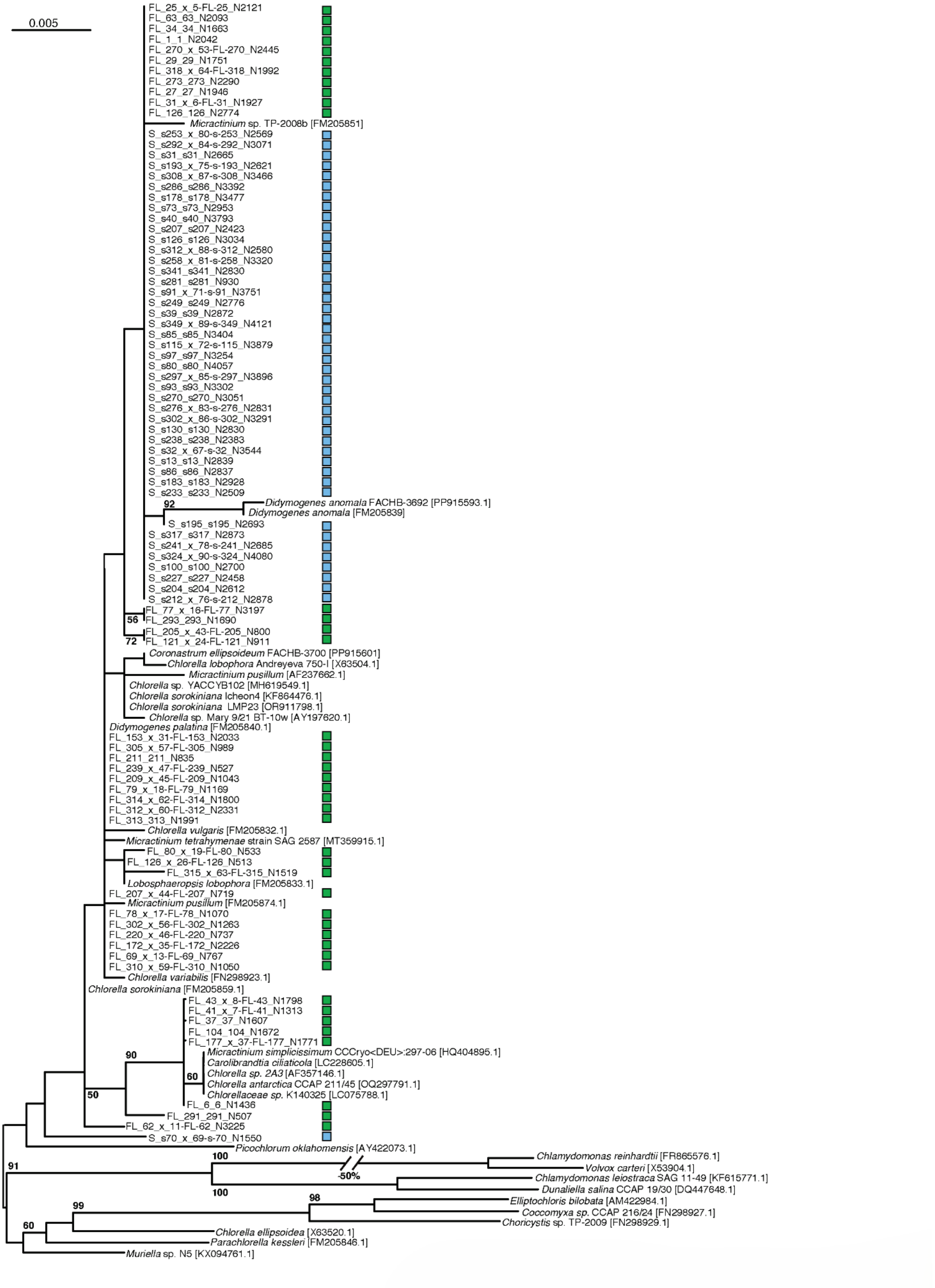


**Fig S1. Phylogeny of 86 strains compared in this project using the front portion of the SSU rRNA encoding gene and demonstrating all strains recovered group closely to known *Micractinium* and *Chlorella* strains.** We note that 41 free-living strains sampled show a higher degree of rDNA variation than the 45 symbiotically isolated strains. The phylogeny was calculated from a masked alignment of 121 sequences and 777 characters using PhyML (REF) with GTR + Γ (with 4 rate categories [α shape parameter = 0.67] 0 and proportion of invariant sites (I = 0.57) correction. 100 bootstrap replicates were conducted.

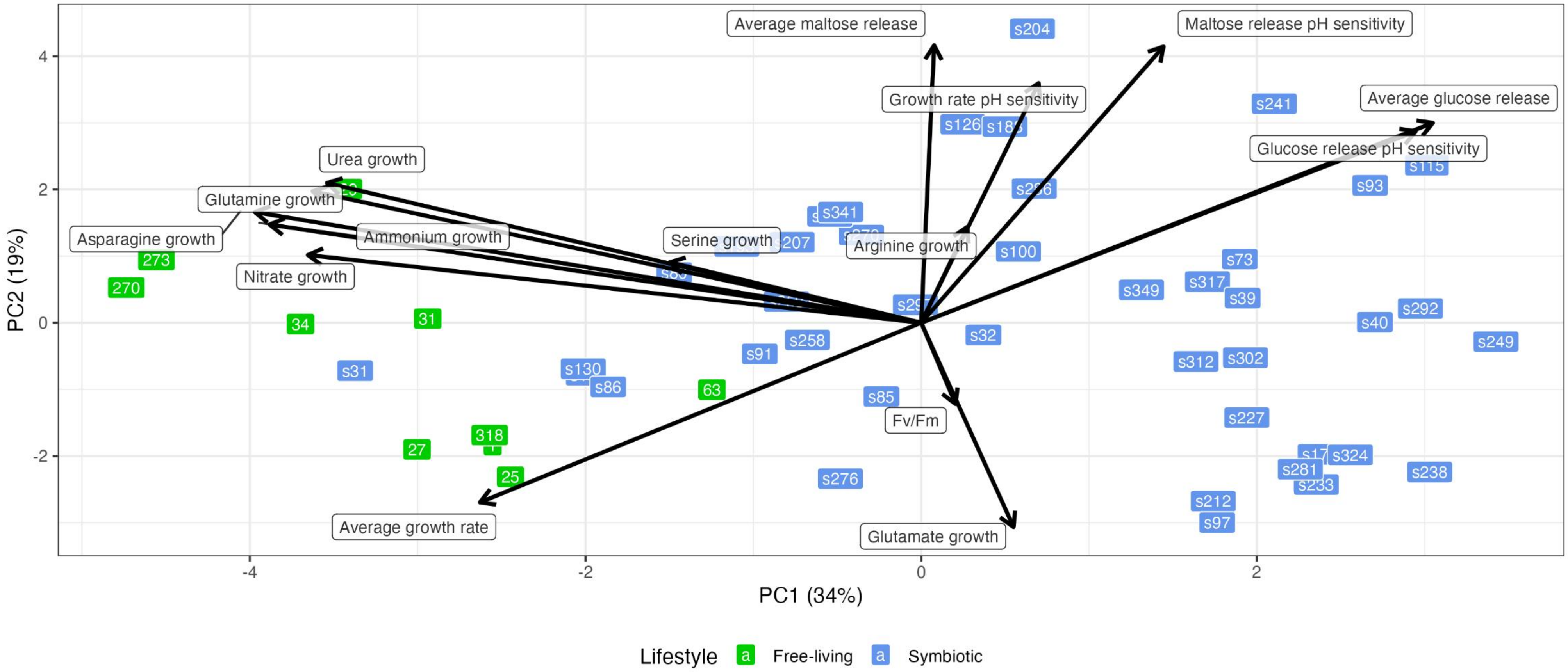


**Fig S2. Phenotypic divergence of endosymbiotic (blue) and free-living (green) strains belonging to phylogroup 1 in multivariate space with variable loadings.** Panels show biplots of principal components from a principal component analysis of multivariate trait data. Arrows and labels show loadings of individual variables. Oblongs indicate the position of individual strains in multivariate trait space and are labelled with the strain number. All strains belong to phylogroup 1 as defined in Fig S1.

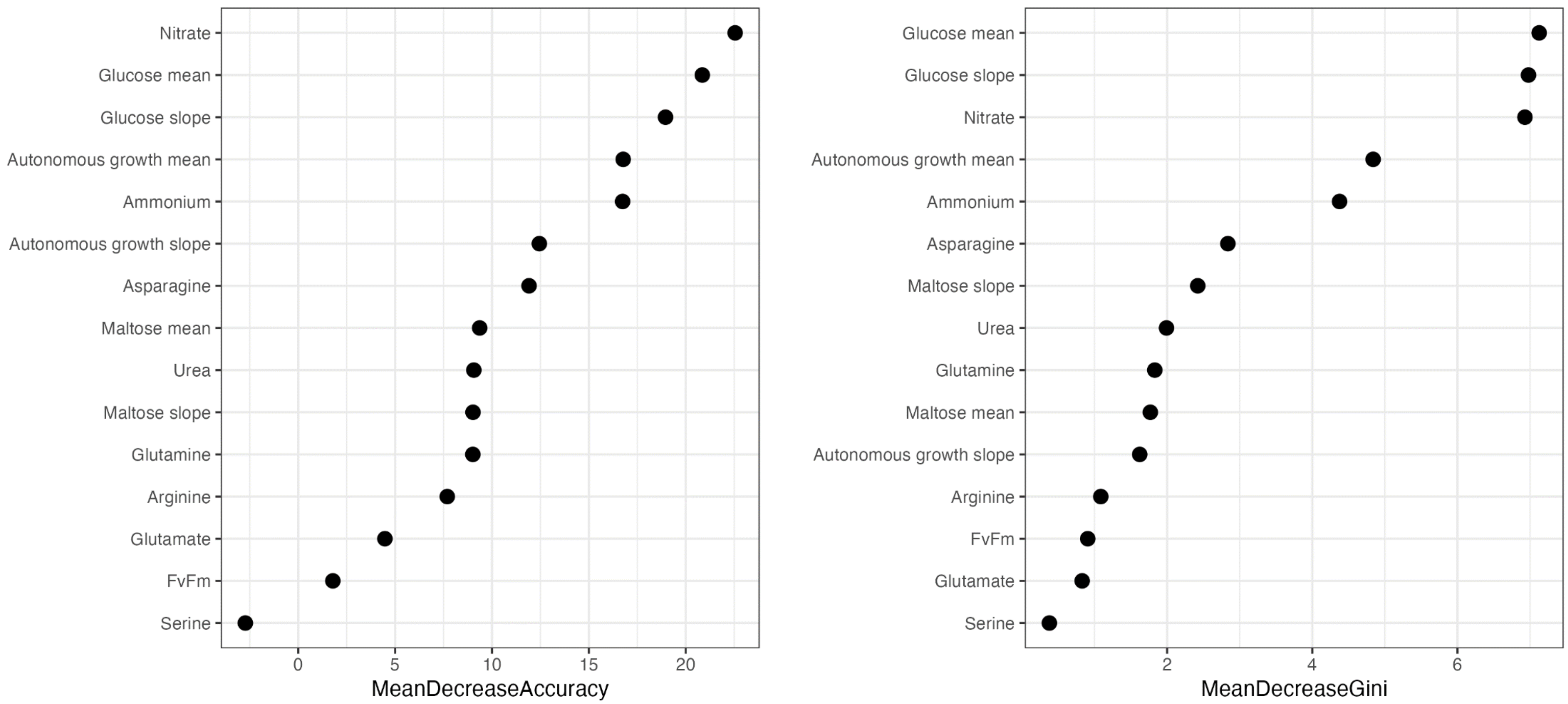


**Fig S3. Phenotypic variables ranked by Mean Decrease Accuracy (importance as predictor) and Mean Decrease Gini (importance for model to be able to make a clean split) for the Random Forest classifier model predicting strain lifestyle.** Where the trait was tested in both neutral and acidic pH, the trait has been summarised as a mean for overall performance and a slope (difference) between performance in the two pHs to indicate pH sensitivity.

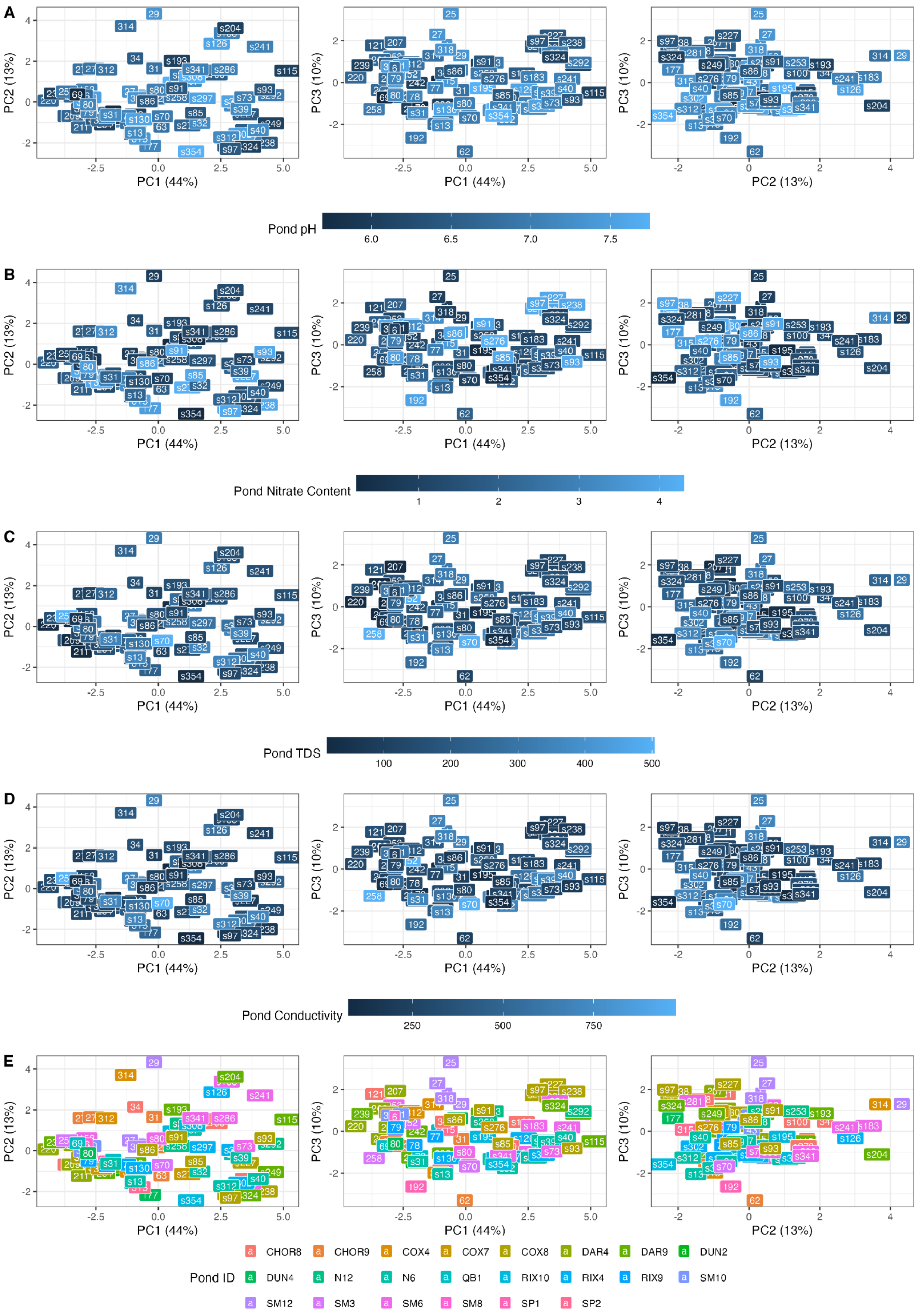


**Fig S4. Principal Component Analysis biplots of multivariate trait data overlayed with environmental variables** a) pond pH, b) pond nitrate content, c) pond TDS, d) pond conductivity and e) pond ID.

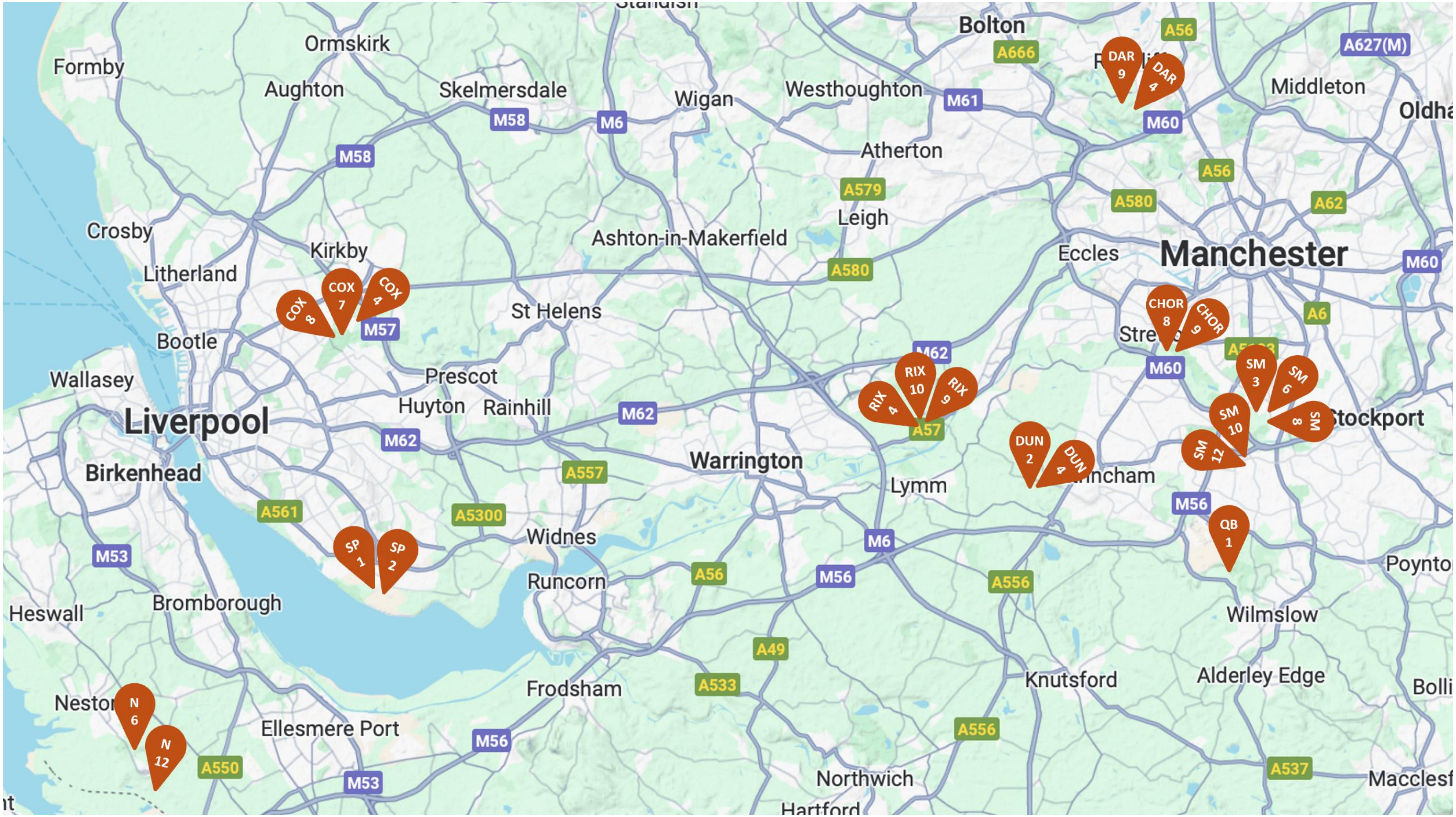


**Fig S5. Map of pond sites in Greater Manchester area, June–September in 2021 and 2022 (Google Maps [created on 24 July 2025]).**
