## Supplementary Tables for "Intracellular niche specialisation drives evolutionary entrapment of endosymbiotic algae"

**Table S1. Model summary output and selected pairwise contrasts from Bayesian hurdle gamma regression model estimating likelihood of sugar release (hurdle) and amount of sugar released when non-zero (mean).**

| Response | Submodel | Term | Estimate | 95% CI (lower–upper) |
|---|---|---|---|---|
| **Glucose** | mean | Intercept | 5.44 | 1.56–9.29 |
| | | pH | −0.73 | −1.43–−0.02 |
| | | lifestyleS | −0.44 | −4.23–3.35 |
| | | pH × lifestyleS | 0.22 | −0.48–0.91 |
| | hurdle | Intercept | −12.03 | −16.38–−7.67 |
| | | pH | 2.66 | 1.87–3.44 |
| | | lifestyleS | −0.86 | −4.60–2.87 |
| | | pH × lifestyleS | −0.69 | −1.37–0.00 |
| **Maltose** | mean | Intercept | 5.52 | 4.62–6.41 |
| | | pH | −0.55 | −0.68–−0.41 |
| | | lifestyleS | 1.99 | 0.90–3.13 |
| | | pH × lifestyleS | −0.26 | −0.43–−0.08 |
| | hurdle | Intercept | −3.73 | −5.27–−2.22 |
| | | pH | 0.68 | 0.45–0.93 |
| | | lifestyleS | −5.57 | −7.90–−3.38 |
| | | pH × lifestyleS | 0.61 | 0.28–0.96 |
| **Trehalose** | mean | Intercept | 1.43 | −2.96–6.29 |
| | | pH | 0.05 | −0.72–0.76 |
| | | lifestyleS | 1.06 | −2.58–4.59 |
| | | pH × lifestyleS | −0.20 | −0.81–0.44 |
| | hurdle | Intercept | 0.27 | −4.05–4.32 |
| | | pH | 0.55 | −0.08–1.27 |
| | | lifestyleS | −0.38 | −3.84–3.11 |
| | | pH × lifestyleS | 0.09 | −0.50–0.68 |
| **Random effects (SD of intercept by strain)** | | glucose | 0.28 | 0.06–0.48 |
| | | maltose | 0.87 | 0.70–1.07 |
| | | trehalose | 0.56 | 0.02–1.52 |
| **Shape parameters** | | glucose | 3.65 | 2.73–4.77 |
| | | maltose | 2.95 | 2.42–3.53 |
| | | trehalose | 1.54 | 0.56–3.65 |

**Selected pairwise contrasts**

| Sugar | Model | Contrast | Estimate | HPD (lower—upper) |
|---|---|---|---|---|
| Glucose | mean | FL—S | -0.26 | -0.95—0.44 |
| | hurdle | FL—S | 0.69 | 0.0056—1.37 |
| Maltose | mean | FL—S | 0.26 | 0.089—0.43 |
| | hurdle | FL—S | -0.61 | -0.93—-0.26 |
| Trehalose | mean | FL—S | 0.21 | -0.44—0.81 |
| | hurdle | FL—S | -0.088 | -0.65—0.52 |

**Table S2. Output from mixed effects model testing effect of lifestyle and N source on growth rate along with selected pairwise contrasts.**

| **Fixed effect** | **X2** | **DF** | **p-value** |
|---|---|---|---|
| Lifestyle | 82.73 | 1 | < 0.001 |
| N source | 722.12 | 7 | < 0.001 |
| Lifestyle:N source | 564.56 | 7 | < 0.001 |
| **Selected pairwise contrasts** | | | |
| **Contrast (FL—S)** | **t-ratio** | **DF** | **p-value** |
| Ammonium | 12.34 | 224 | < 0.001 |
| Arginine | 3.08 | 224 | 0.15 |
| Asparagine | 12.38 | 224 | < 0.001 |
| Glutamate | -2.91 | 224 | 0.22 |
| Glutamine | 12.39 | 224 | < 0.001 |
| Nitrate | 11.12 | 224 | < 0.001 |
| Serine | 2.50 | 224 | 0.48 |
| Urea | 6.99 | 224 | < 0.001 |

**Table S3. Output from envfit analysis showing correlation of phenotypic variables with PCA axes.** The trait was tested in both neutral and acidic pH and has been summarised as a mean for overall performance and a slope (difference) between performance in the two pHs to indicate pH sensitivity.

| Variable | PC1 | PC2 | $r^2$ | p-value |
|---|---|---|---|---|
| Glucose (mean) | 0.796 | 0.606 | 0.815 | 0.001 |
| Maltose (mean) | 0.374 | 0.927 | 0.805 | 0.001 |
| Glucose (slope) | 0.8 | 0.6 | 0.781 | 0.001 |
| Maltose (slope) | 0.466 | 0.885 | 0.842 | 0.001 |
| Arginine | −1.000 | 0.002 | 0.186 | 0.001 |
| Asparagine | −0.718 | 0.696 | 0.818 | 0.001 |
| Glutamine | −0.641 | 0.768 | 0.77 | 0.001 |
| Glutamate | 0.865 | 0.502 | 0.144 | 0.001 |
| Serine | −0.726 | 0.688 | 0.137 | 0.002 |
| Ammonium | −0.871 | 0.491 | 0.868 | 0.001 |
| Nitrate | −0.842 | 0.54 | 0.766 | 0.001 |
| Urea | −0.758 | 0.653 | 0.775 | 0.001 |
| Autonomous growth (mean) | −1.000 | −0.023 | 0.495 | 0.001 |
| Autonomous growth (slope) | 0.849 | 0.528 | 0.324 | 0.001 |
| Fv/Fm | 0.836 | 0.549 | 0.065 | 0.054 |

**Table S4. Results of envfit analysis showing correlations between individual traits and the first two principal components of the multivariate trait ordination.** Vector directions indicate the orientation of increasing trait values in ordination space.

| Trait | PC1 loading | PC2 loading | $r^2$ | P |
|---|---|---|---|---|
| Glucose (mean) | 0.796 | 0.606 | 0.815 | 0.001 |
| Maltose (mean) | 0.374 | 0.927 | 0.805 | 0.001 |
| Glucose (slope) | 0.8 | 0.6 | 0.781 | 0.001 |
| Maltose (slope) | 0.466 | 0.885 | 0.842 | 0.001 |
| Arginine | −1.000 | 0.002 | 0.186 | 0.001 |
| Asparagine | −0.718 | 0.696 | 0.818 | 0.001 |
| Glutamine | −0.641 | 0.768 | 0.77 | 0.001 |
| Glutamate | 0.865 | 0.502 | 0.144 | 0.002 |
| Serine | −0.726 | 0.688 | 0.137 | 0.002 |
| Ammonium | −0.871 | 0.491 | 0.868 | 0.001 |
| Nitrate | −0.842 | 0.54 | 0.766 | 0.001 |
| Urea | −0.758 | 0.653 | 0.775 | 0.001 |
| Autonomous growth (mean) | −1.000 | −0.023 | 0.495 | 0.001 |
| Autonomous growth (slope) | 0.849 | 0.528 | 0.324 | 0.001 |
| Fv/Fm | 0.836 | 0.549 | 0.065 | 0.048 |

**Table S5. PERMANOVA output testing effects on multivariate centroids of lifestyle interacting with pond identity and associated environmental variables, as well as betadisper output for homogeneity of dispersion tests.** Pond identity and environmental variables were tested in separate models.

| Model / Term | Effect | Df | $R^2$ | F | P | Dispersion F | Dispersion P |
|---|---|---|---|---|---|---|---|
| Lifestyle only | Lifestyle | 1, 91 | 0.311 | 41.01 | 0.001 | 1.88 | 0.173 |
| Lifestyle × Pond ID | Lifestyle | 1, 65 | 0.311 | 49.5 | 0.001 | – | – |
| | Pond ID | 21, 65 | 0.25 | 1.9 | 0.001 | 3.41 | <0.001 |
| | Lifestyle × Pond ID | 5, 65 | 0.032 | 1.01 | 0.426 | – | – |
| Lifestyle × Nitrate | Lifestyle | 1, 89 | 0.311 | 42.39 | 0.001 | – | – |
| | Nitrate | 1, 89 | 0.019 | 2.57 | 0.048 | 3.14 | <0.001 |
| | Lifestyle × Nitrate | 1, 89 | 0.018 | 2.49 | 0.046 | – | – |
| Lifestyle × TDS | Lifestyle | 1, 89 | 0.311 | 41.65 | 0.001 | – | – |
| | TDS | 1, 89 | 0.008 | 1.05 | 0.341 | 3.51 | <0.001* |
| | Lifestyle × TDS | 1, 89 | 0.018 | 2.36 | 0.05 | – | – |
| Lifestyle × pH | Lifestyle | 1, 89 | 0.311 | 42.92 | 0.001 | – | – |
| | pH | 1, 89 | 0.012 | 1.67 | 0.127 | 3.45 | <0.001 |
| | Lifestyle × pH | 1, 89 | 0.033 | 4.57 | 0.004 | – | – |
| Lifestyle × Conductivity | Lifestyle | 1, 89 | 0.311 | 41.85 | 0.001 | – | – |
| | Conductivity | 1, 89 | 0.008 | 1.05 | 0.317 | 3.41 | <0.001 |
| | Lifestyle × Conductivity | 1, 89 | 0.021 | 2.82 | 0.032 | – | – |

**Table S6. Site details, mean values of physicochemical parameters measured *in situ* June–September in 2021 and 2022.**

| Pond site | Site name | OS Grid Reference | pH | Conductivity, µS/cm | TDS, ppm | $NO^3$, mg/L |
|---|---|---|---|---|---|---|
| | | | Mean (±SD) | | | |
| CHOR8 | Chorlton Ees and Ivy green LNR | SJ 80405 93225 | 6.56 (0.22) | 284 (83) | 141 (40) | 0.85 (0.65) |
| CHOR9 | | SJ 80355 93095 | 6.77 (0.33) | 246 (33) | 127 (17) | 1.41 (1.20) |
| COX4 | Croxteth Hall and Country Park | SJ 41975 94725 | 6.84 (0.14) | 410 (106) | 242 (115) | 3.33 (0.65) |
| COX7 | | SJ 40974 94870 | 6.66 (0.22) | 310 (191) | 138 (58) | 4.30 (2.20) |
| COX8 | | SJ 40900 94858 | 6.43 (0.32) | 190 (35) | 128 (73) | 4.06 (1.02) |
| DAR4 | Darcy Lever gravel pits, Moses Gate Country Park | SD 74245 07745 | 6.84 (0.26) | 381 (119) | 124 (154) | 1.94 (0.00) |
| DAR9 | | SD 74115 07845 | 6.26 (0.57) | 197 (165) | 100 (84) | 1.30 (0.00) |
| DUN2 | Dunham Massey Par (NT), Oxbow lakes | SJ 73365 86975 | 7.13 (0.20) | 363 (55) | 181 (27) | 1.62 (1.86) |
| DUN4 | Dunham Massey Park (NT) | SJ 73715 87125 | 7.12 (0.11) | 453 (99) | 213 (26) | 2.26 (2.35) |
| N6 | Ness, The University of Liverpool Botanical Gardens | SJ 30445 75355 | 7.32 (0.32) | 633 (133) | 320 (62) | 2.12 (0.01) |
| N12 | Burton Mere Wetlands (RSPB), Dee Estuary Nature Reserve | SJ 31395 73565 | 7.00 (0.10) | 429 (76) | 239 (54) | 1.35 (0.05) |
| QB1 | Quarry Bank (NT), upper sheep fields | SJ 82395 83125 | 5.85 (0.27) | 105 (33) | 53 (14) | 1.20 (0.96) |
| RIX4 | Rixton Claypits (LNR) | SJ 68585 90515 | 7.70 (0.24) | 601 (27) | 300 (13) | 0.98 (0.48) |
| RIX9 | Rixton, Burton Mere House (RSPB) | SJ 68705 90635 | 7.09 (0.26) | 480 (238) | 185 (69) | 2.00 (2.56) |
| RIX10 | Rixton Claypits (LNR) | SJ 68505 90685 | 8.45 (1.25) | 102 (38) | 51 (19) | 0.44 (0.29) |
| SM3 | South Manchester, Didsbury rock garden / Fletcher Moss Botanical Gardens, north site | SJ 84537 90383 | 7.05 (0.09) | 992 (23) | 504 (1) | 1.00 (0.00) |
| SM6 | South Manchester, Didsbury rock garden / Fletcher Moss Botanical Gardens | SJ 84763 90287 | 7.20 (0.23) | 345 (125) | 192 (46) | 0.93 (0.28) |
| SM8 | South Manchester, Fletcher Moss Park | SJ 84724 90074 | 6.91 (0.31) | 265 (65) | 131 (31) | 1.00 (0.00) |
| SM10 | South Manchester, Gatley Carrs (WT) | SJ 84075 88915 | 7.55 (0.15) | 595 (44) | 305 (32) | 0.60 (0.25) |
| SM12 | | SJ 84045 88835 | 7.43 (0.19) | 653 (144) | 304 (35) | 0.63 (0.35) |
| SP1 | Speke Hall (NT) | SJ 41908 82709 | 6.97 (0.11) | 533 (18) | 266 (6) | 3.92 (2.38) |
| SP2 | | SJ 41918 82725 | 7.42 (0.02) | 557 (0) | 279 (0) | 3.71 (0.00) |

**Table S7. Alterations of MBBM medium made for algal biomass growth and experiment of each assay.**

<table>
<tr><th colspan="2" rowspan="2">Experiment</th><th colspan="2">N source*</th><th rowspan="2">Sucrose*</th><th colspan="2">pH**</th></tr>
<tr><th>$NaNO_3$</th><th>Peptone</th><th>Value</th><th>Adjusted with</th></tr>
<tr><td rowspan="9">Growth rate on different organic and inorganic nitrogen (N) sources</td><td>To grow biomass</td><td>Yes</td><td>Yes</td><td>Yes</td><td>7.6</td><td>NaOH</td></tr>
<tr><td rowspan="8">For experiment</td><td colspan="2">Sodium nitrate - 0.38 g/L</td><td rowspan="8">Yes</td><td rowspan="8">7.6</td><td rowspan="8">NaOH, HCl</td></tr>
<tr><td colspan="2">Ammonium chloride - 0.58 g/L</td></tr>
<tr><td colspan="2">Urea - 0.64 g/L</td></tr>
<tr><td colspan="2">L-Arginine - 1.30 g/L</td></tr>
<tr><td colspan="2">L-Glutamine - 0.93 g/L</td></tr>
<tr><td colspan="2">L-Glutamate (L-Glutamic acid) - 0.75 g/L</td></tr>
<tr><td colspan="2">L-Asparagine - 0.47 g/L</td></tr>
<tr><td colspan="2">L-Serine - 0.53 g/L</td></tr>
<tr><td rowspan="2">Growth rate in acidic and neutral medium</td><td>To grow biomass</td><td rowspan="2">Yes</td><td rowspan="2">Yes</td><td rowspan="2">No</td><td>7.6</td><td>NaOH</td></tr>
<tr><td>For experiment</td><td>5.5 and 7.6</td><td>MES monohydrate and MES Na salt</td></tr>
<tr><td rowspan="2">Screening of sugar release</td><td>To grow biomass</td><td>Yes</td><td>Yes</td><td rowspan="2">No</td><td>7.6</td><td>NaOH</td></tr>
<tr><td>For experiment</td><td>No</td><td>No</td><td>5.5 and 7.6</td><td>MES monohydrate and MES Na salt</td></tr>
<tr><td rowspan="2">Photosynthetic efficiency</td><td>To grow biomass</td><td rowspan="2">Yes</td><td rowspan="2">Yes</td><td rowspan="2">No</td><td rowspan="2">7.6</td><td rowspan="2">NaOH</td></tr>
<tr><td>For experiment</td></tr>
</table>

* - Standard elements of MBBM

** - Standard pH of MBBM around 6.6 (not adjusted)

**Table S8. Methods of Multi-Step Gradient for sugar release detection in different pH media.**

**Method for samples in pH 5.5 medium**

| | |
|---|---|
| A, % – 30 mM NaOH<br>B, % – 100 mM NaOH<br>C, % – 300 mM NaOH-500 mM NaOAc<br>D, % – MiliQ | **Retention time:**<br>Trehalose – 2.2 ± 2 min<br>Glucose – 4.7 ± 2 min<br>Maltose – 19.8 ± 2 min |

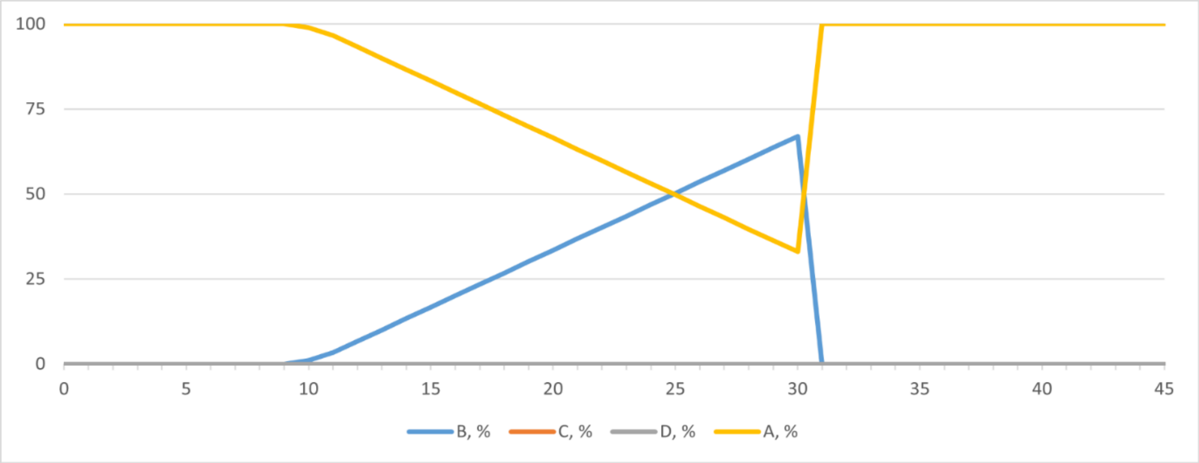


| Time, min | Flow, ml/min | B, % | C, % | D, % | Curve |
|---|---|---|---|---|---|
| 0.00 | Run | | | | |
| 0.00 | 0.4 | 0.0 | 0.0 | 0.0 | 5 |
| 10.00 | 0.4 | 0.0 | 0.0 | 0.0 | 5 |
| 30.00 | 0.4 | 67.0 | 0.0 | 0.0 | 5 |
| 31.00 | 0.4 | 0.0 | 0.0 | 0.0 | 5 |
| 45.00 | 0.4 | 0.0 | 0.0 | 0.0 | 5 |
| 45.00 | Stop Run | | | | |

**Method for samples in pH 7.6 medium**

| | |
|---|---|
| A, % – 30 mM NaOH<br>B, % – 30 mM NaOH-500 mM NaOAc<br>C, % – 300 mM NaOH-500 mM NaOAc<br>D, % – MiliQ | **Retention time:**<br>Trehalose – 2.5 ± 2 min<br>Glucose – 7.5 ± 2 min<br>Maltose – 20.0 ± 2 min |

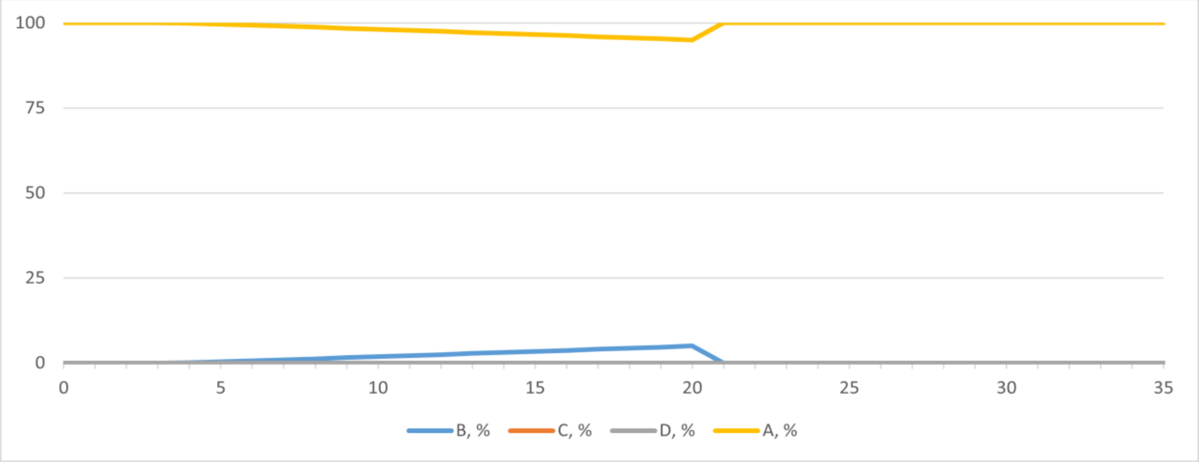


| Time, min | Flow, ml/min | B, % | C, % | D, % | Curve |
|---|---|---|---|---|---|
| 0.00 | Run | | | | |
| 0.00 | 0.4 | 0.0 | 0.0 | 0.0 | 5 |
| 4.00 | 0.4 | 0.0 | 0.0 | 0.0 | 5 |
| 20.00 | 0.4 | 5.0 | 0.0 | 0.0 | 5 |
| 20.50 | 0.4 | 0.0 | 0.0 | 0.0 | 5 |
| 35.00 | 0.4 | 0.0 | 0.0 | 0.0 | 5 |
| 35.00 | Stop Run | | | | |
